# Two opposing effects of monovalent cations on the stability of i-motif structure

**DOI:** 10.1101/2022.09.30.510309

**Authors:** Sung Eun Kim, Seok-Cheol Hong

## Abstract

At acidic pH, cytosine-rich single-stranded DNA can be folded into a tetraplex structure called i-motif (iM). In recent studies, the effect of monovalent cations on the stability of iM structures has been addressed, but a consensus about the issue has not been reached yet. Thus, we investigated the effects of various factors on the stability of iM structures using fluorescence resonance energy transfer (FRET)-based analysis for three types of iM derived from human telomere sequences. We confirmed that the iM structure is destabilized as the concentration of monovalent cations (Li^+^, Na^+^, K^+^) increases and that Li^+^ has the greatest tendency of destabilization. This cation-induced destabilization is rather unexpected and specific to the iM structure, considering the cation’s electrostatic effect of supporting DNA folding. Monovalent cations of different kinds promote the flexibility of single-stranded DNA (ssDNA) and the stability of folded DNA structures to different degrees, suggesting that the size of cation be a key factor in its function. All taken together, we conclude that the stability of iM structures is controlled by the subtle balance of the two counteractive effects of monovalent cations, electrostatic screening and disruption of cytosine base pairing.

## I. INTRODUCTION

Maintenance of telomere regions is crucial for cell survival. The human telomere regions contain a special repeat sequence (TAACCC), which is thought to play an important role in protecting their termini from nucleases.^1,2^ The cytosine-rich sequence was found to form a folded quadruplex structure named i-motif (iM) (Figure 1a).^3^ The iM structure occurs in acidic conditions because C:C^+^ base pairing is required for iM formation and a protonated cytosine(C^+^) can be formed more easily in acidic conditions (Figure 1b).^4–6^ Although there have been doubts about the presence of iM structures in living cells with a physiological pH, several studies have shown that iM structures may be stabilized in crowded environments at neutral pH^7–9^ and the structures are formed in the nuclei of human cells.^10,11^ Although the biological function of iM is still under debate, this pH-sensitive molecular motif holds great promise in the fields of cell biology and nanotechnology.^12–16^

**Figure 1.**
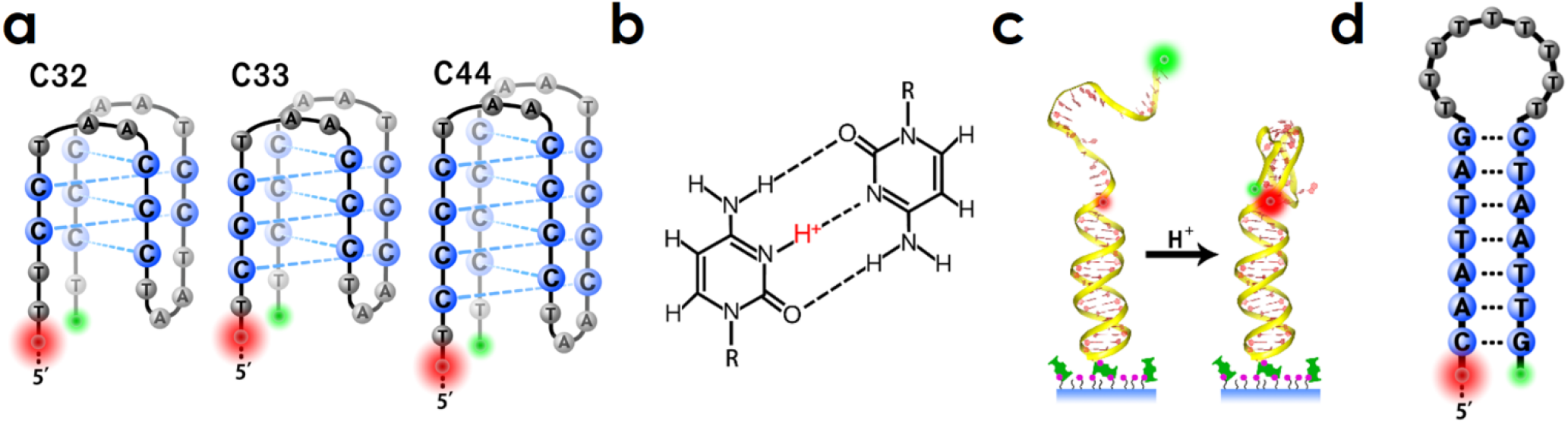
(a) Three types of iM molecules used in this work. The sequence of iM molecules and the positions of Cy3 and Cy5 dyes for FRET assays are also shown. Dotted lines indicate C:C^+^ base pairing. (b) The base pair between cytosine and protonated cytosine (C:C^+^ pair) (c) Schematic picture of folding of the iM structure at high [H^+^] (d) Hairpin DNA molecule used in this work. The sequence of hairpin DNA and the positions of Cy3 and Cy5 dyes for FRET assay are shown. In (a) and (d), bases in loops are drawn smaller and in gray.

In our previous study, we investigated the effect of monovalent cations on the stability of iM structures using single-molecule FRET (fluorescence resonance energy transfer) technique because the technique is ideal for detecting conformational transitions occurring at nanoscale. In the study, we found that the iM structure becomes unstable with increasing concentration of Li^+^.^17^ Two recent works addressed the effect of monovalent cations on iM. Zhang et. al, reported that monovalent cations such as Li^+^, Na^+^, and K^+^ have a destabilizing effect on the iM structure while Gao et. al, argued that the effect of K^+^ is rather puzzling: it either stabilizes or destabilizes iM, depending on the pH and type of buffer.^18,19^ Such an inconsistency and ambiguity in this field motivated us to revisit this problem in a more systematic manner. Here, we examined the stability of three types of iM DNA using bulk and single-molecule FRET techniques. Considering the universal stabilizing effect of monovalent cations on a hairpin DNA, the main structural motif of which is dsDNA, we shall conclude that not only Li^+^ but also Na^+^ and K^+^ have a destabilizing effect on the iM structure aside from their well-known electrostatic stabilization. Interestingly, Li^+^ facilitates the formation of iM more than Na^+^ or K^+^ at the lower range of concentration tested (< 200 mM). This implies that Li^+^ is more effective than other monovalent cations in the formation of iM although Li^+^ strongly destabilizes the same structure at higher concentrations (> 200 mM). This rather complex effect of cations on folded DNA structures can be accounted for by the two counteractive actions of monovalent cations, (i) reduced electrostatic repulsion, which also makes ssDNA flexible for easier folding, and (ii) disruption of hydrogen bonds between cytosine bases. In both effects, Li^+^ is distinctive likely because of its smaller size.

## II. MATERIALS AND METHODS

### 2.1. Samples

In this study, three different cytosine-rich oligonucleotides (C32, C33, and C44; see Table 1 and Figure 1a) were used to form iM structures. C32, C33, and C44 have up to 5, 6, and 8 C:C^+^ base pairs when they are intramolecularly folded. C33 has the same sequence as the C-rich human telomere sequence: there are four cytosine triplets, which are separated by a non-cytosine triplet (TAA). In C32, the first cytosines of the first and third cytosine triplets are replaced with thymines so that the molecule once folded has one less C:C^+^ base pairs. In C44, all cytosine triplets are replaced with cytosine quartets so that the molecule once folded has two more C:C^+^ base pairs than C33. The underlined part of the sequences forms the iM structure in Table 1. These oligonucleotides are labeled with Cy3 and Cy5 fluorescent dyes for FRET assays as shown in Figure 1a. In bulk FRET assays, each iM oligo was used as described below. To perform smFRET measurements, each iM oligo was hybridized with the stem oligo (see Table 1). The stem oligo has a biotin at its 3’-end, which allows the assembled DNA to bind to the NeutrAvidin-coated glass substrate. The thermal stability of a hairpin DNA was also tested with bulk FRET assays in comparison with iM DNA. All oligos were purchased from IDT (Integrated DNA Technologies, USA).

**Table 1.**
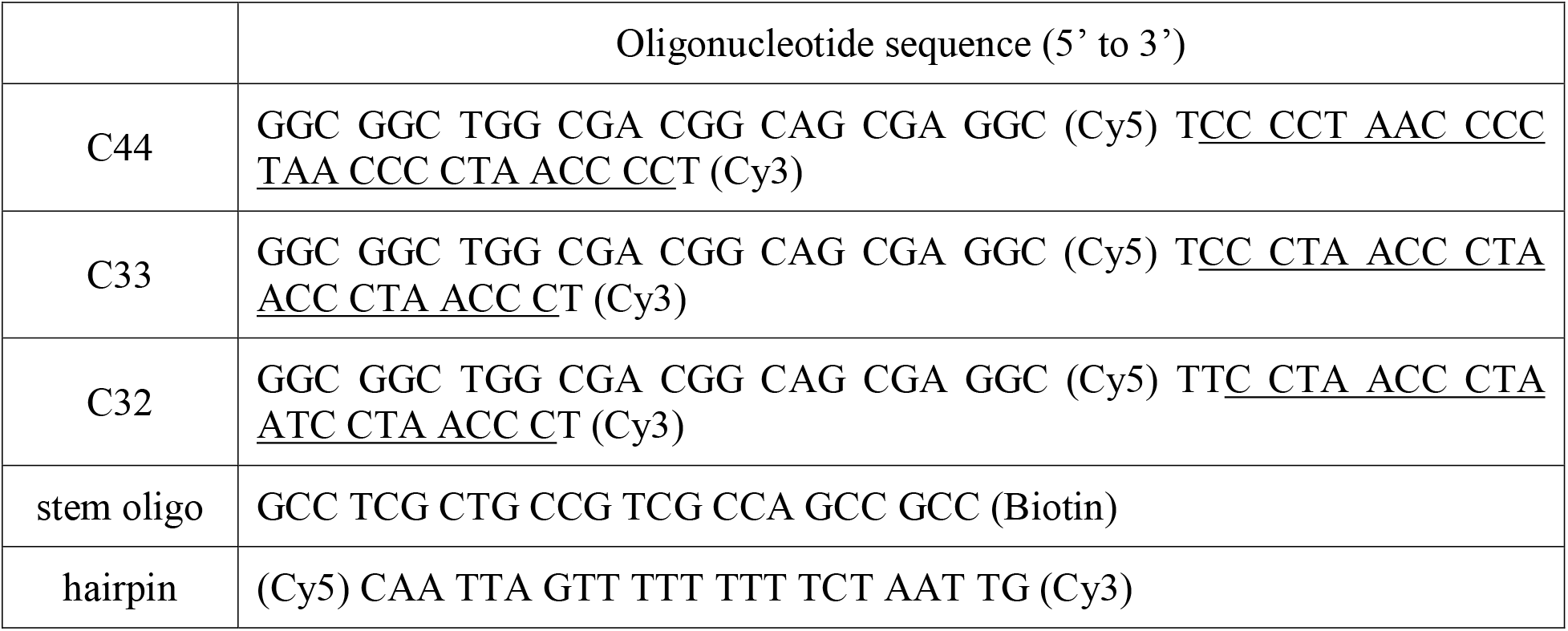
Oligonucleotide sequences for a hairpin DNA and three different types of i-motif DNA together with a stem strand.

### 2.2. Bulk FRET assay

Fluorescence spectra were acquired using an FS-2 fluorescence spectrometer (Scinco, Korea) equipped with a Peltier temperature controller. We measured fluorescence spectra of iM samples (100 nM) in 20 mM sodium acetate buffer with pH 4.5, 5.0, and 5.5, containing various concentrations of cations (50~500 mM). The pH of this buffer remains unchanged over a wide range of temperature, so this buffer is ideal for the melting assays of iM.^20,21^ Since bulk FRET assay does not require immobilization of DNA molecules to the bottom of a sample chamber, this assay was performed with one of the iM oligos alone without the stem oligo. We performed this assay and acquired fluorescence spectra as follows: iM molecules loaded in a standard quartz cuvette were excited at 532 nm and their emission spectra were measured from 550 nm to 690 nm with the scanning interval and speed of 0.1 nm and 60 nm/min, respectively. The melting assay via FRET measurement was carried out by increasing the temperature from 19°C to 85~91°C in 3-degree increments and measuring spectra at each target temperature after reaching it and staying there stably for 5 minutes. The average emission intensity of Cy3 over the range of 555-565 nm was normalized to 1 and the average Cy5 emission intensity over the range of 650-680 nm was assumed to be proportional to the number of folded molecules. With increasing temperature, the fluorescence intensities of Cy3-only or Cy5-only samples decreases. To compensate for this temperature dependent intensity change, we also performed control experiments with Cy3-only and Cy5-only oligos to measure the effect of temperature on Cy3 and Cy5 emission. Using dyes’ temperature characteristics, bulk FRET data were corrected (Figure S1, S3).

### 2.3. Single-molecule FRET assay

Single-molecule FRET (smFRET) experiments were performed using a TIRF (Total Internal Reflection Fluorescence) microscope (Nikon ECLIPSE Ti-U). The stem oligo containing a biotin was hybridized with an iM oligo and then attached to the NeutrAvidin-coated glass. T50 buffer (10 mM Tris-HCl, 50 mM NaCl, pH 7.4) containing a low concentration (~25 pM) of DNA was injected into the chamber to fix the DNA sample on the glass. Then we washed out the chamber with 400 μL imaging buffer, which consisted of 380 μL of pH buffer (50 mM MES) containing 2 mM Trolox (Sigma Aldrich) supplemented with 16 μL of 10%(w/v) D-(+)-glucose (Sigma-Aldrich) and 4 μL of Gloxy (100×). The Gloxy stock solution (100×) was prepared by mixing 0.04 g of glucose oxidase (168800 units/g, Sigma-Aldrich), 125 μL of catalase (16 mg protein/mL, 42600 units/mg, Sigma-Aldrich), and 260 μL of T50. Fluorescent dyes are readily oxidized and lose fluorescence, so a measure to remove oxygen from the chamber using an oxygen scavenging system (mixed with glucose and gloxy) is highly recommended. Gluconic acid produced by the oxygen scavenger system, however, lowers the level of pH inside the chamber, so pH buffering is required (in our assay, we use 50 mM MES buffer).^22^ The FRET efficiency (*E*_FRET_) from each pair of dyes (donor and acceptor) is calculated by *E*_FRET_ = *I_A_*/(*I_A_* + *I_D_*) where *I_D_* and *I_A_* are the fluorescence intensities of donor and acceptor dyes. Single-molecule FRET experiments were performed at room temperature in buffers at various pH and salt conditions.

## III. RESULTS & DISCUSSIONS

### pH dependent formation of the iM structure

A cytosine-rich strand adopts the four-stranded iM structure at low pH conditions, which can be sensitively measured by FRET. With Cy3 (donor) and Cy5 (acceptor) dyes at the opposite termini of the iM molecule, the distance between the two dyes changes as the shape of the molecule changes (Figure 1a). When the iM DNA is folded (iM formation), the energy transfer from Cy3 to Cy5 is facilitated due to reduced inter-dye distance. We confirmed by smFRET measurements that iM molecules are folded at low pH, which is favorable under cytosine protonation, but they are unfolded at high pH. At [Na^+^] = 200 mM, there are only two states: unfolded identified by *E*_FRET_~0.3 and folded by *E*_FRET_~0.9 (Figure S2).

### C:C^+^ base-pair dependent stability of the iM structure

In nature, there are many cytosine-rich sequences with various numbers of cytosines, some of which are known to form the iM structure.^23–28^ In order to understand how the stability of the iM structure depends on the number of C:C+ base pairs, we first performed bulk FRET-based iM melting assays on three different C-rich molecules (C32, C33, and C44) at pH 4.5, 5.0, and 5.5. When an iM molecule is thermally melted and unfolded, the FRET efficiency is indeed decreased. One might think that *I*_D_ and *I*_A_ would exhibit anti-correlation, that is, *I*_D_ increases and *I*_A_ decreases due to the decrease of energy transfer. This is not exactly the case. There is a subtle issue of temperature-dependent fluorescence efficiency of dye molecules. The issue of proper normalization and calibration of fluorescence efficiency at different temperatures is addressed in SI and handled accordingly hereafter (Figure S3). In the bulk FRET assays, the lower the pH value is or the more C:C^+^ base pairs we have, the higher *T_m_* results, indicating more stable iM. At the acidic conditions of pH = 5.5, 5.0, and 4.5, the melting temperature of iM increases with the number of C:C^+^ base pairs, rising 5.6℃, 6.2℃, and 6.8℃ per C:C^+^ pair, respectively (Figure 2b). At lower pH, the slope of the melting curve becomes steeper, indicating that the degree of stabilization by additional C:C^+^ base pair is stronger at lower pH. Previous studies reported that *T_m_* would increase by ~3.7°C per C:C^+^ pair at pH 5.0 from CD experiments.^29^ The apparent discrepancy is, however, mistaken: the increment of melting temperature per C:C^+^ pair was obtained by linearly fitting the nonlinear curve of *T_m_* vs. the number of C:C^+^ pair over the whole range of their data. As the number of C:C^+^ base pairs increases, *T_m_* increases, but the slope decreases with the range of the number of C:C^+^ base pairs: when the number of C:C^+^ base pairs is limited to the range from 5 to 8 and the curve on this range is linearly fitted, the increment of melting temperature per C:C^+^ pair turns out to be 6.6°C per C:C^+^ pair, which is consistent with our experimental results.

**Figure 2.**
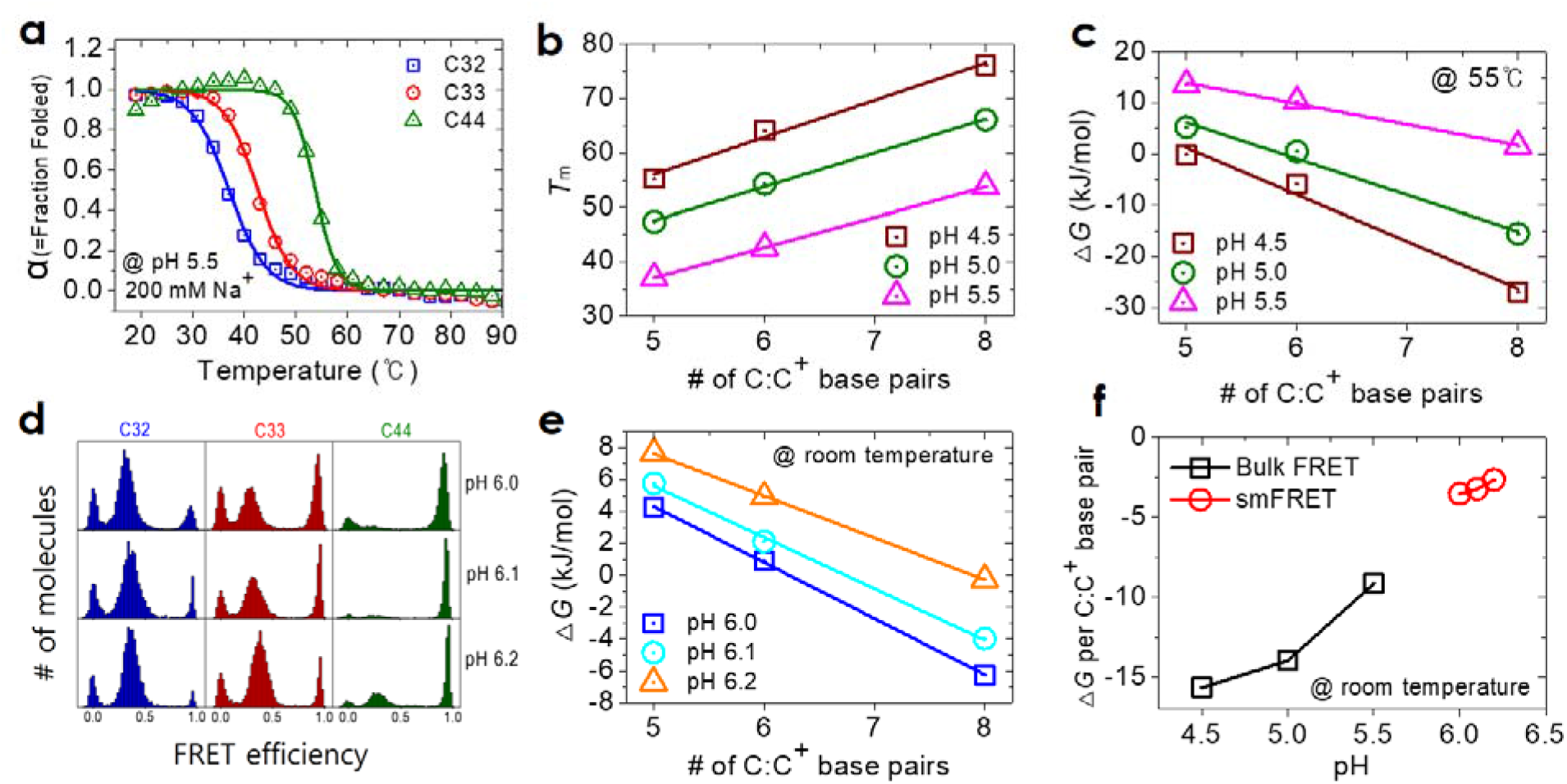
Thermodynamic behavior of iM molecules with various numbers of C:C^+^ base pairs. When folded, C32, C33, and C44 have up to 5, 6, and 8 C:C^+^ base pairs. (a-c, bulk FRET assay) (a) Melting curve of iM structures by C32, C33, and C44 at pH 5.5 and [Na^+^] = 200 mM (b) Melting temperature of iM structures by C32, C33, and C44 at pH 4.5, 5, and 5.5. At each pH, the melting temperature changes linearly with the (maximum) number of C:C^+^ base pairs. (c) Free energy of iM structures by C32, C33, and C44 at pH 4.5, 5, and 5.5. (d-f, smFRET assay) (d) FRET histograms of C32, C33, and C44 molecules at pH 6.0, 6.1, and 6.2 (e) Free energy of iM structures by C32, C33, and C44 at pH 6.0, 6.1, and 6.2 (f) Relation of Δ*G*/(C: C^+^ base pair) versus pH acquired at room temperature using bulk FRET and smFRET. Lines in (b, c, and e) are linear fits to the data obtained for the indicated pH conditions.

We also acquired the folding free energy (Δ*G*) at *T* = 55°C as presented in Figure 2c: it is calculated by Δ*G* = −*RT*ln*K*_eq_ where the equilibrium constant is defined by *K*_eq_ = [folded]/[unfolded] = α/(1-α) with α the fraction of folded state. As expected, the folding free energy becomes more negative with increase of acidity and exhibits the linear change with the number of C:C^+^ base pairs.

It is of practical significance to understand the stability of iM at higher pH. At higher pH, the fraction of iM would be significantly reduced and bulk FRET assays turn out to be insensitive for such a minor population of iM. Thus, we switched to smFRET because it is considerably more sensitive than bulk FRET. C32, C33, and C44 molecules were tested at various pH values (6.0, 6.1, and 6.2) at room temperature at [Na^+^] = 200 mM. Three peaks appear in the FRET histogram (Figure 2d). The peak at zero FRET efficiency represents donor-only molecules whose acceptor dye has been bleached out or cannot be fluorescent for any reason. This peak shall be ignored. The peak at the middle FRET values between 0.2 and 0.4 represent unfolded molecules. The peak at the high FRET value of approximately 0.9 represent iM molecules. The folding free energy (Δ*G*) here is similarly calculated by Δ*G* = −*RT*ln*K*_eq_ where *K*_eq_ = [folded]/[unfolded]=[high FRET area]/[low FRET area]. As expected, a iM molecule with more C:C^+^ base pairs has a lower free energy and maintains a stable iM state (Figure 2c,e). It is notable that Δ*G* per base pair, slope of the graphs in Figure 2c,e, becomes more negative as the value of pH decreases (Figure 2f). This indicates easier protonation of cytosine and stronger binding of each C:C^+^ base pair at lower pH conditions. Here, the free energy per base pair at the low pH values (4.5 – 5.5) was converted to the value at room temperature by extrapolating the results obtained with bulk FRET assays conducted at higher temperatures. In this free energy measurement, the two different methods, bulk and single-molecule FRET, covered separate pH ranges, lower (pH = 4.5 to 5.5) and higher (pH = 6.0 to 6.2), respectively, and over the entire range of pH, Δ*G* per base pair is nearly linearly correlated with the pH value, indicating that the two approaches yield a consistent trend.

Our observation also reveals a general feature of phase transition: as shown in Figure 2a, the melting transition becomes sharper when the size of a system is larger. C44 molecules exhibit the melting transition over ~10°C while C32 molecules undergo the transition over a wider range of ~25°C. The range of temperature over which the transition takes place decreases as the number of C:C^+^ base pairs increases. (Figure S4).

### Salt Dependence of iM Formation

Nucleic acids take their native structures and play biological functions when they are in physiological ionic conditions. iM molecules are not an exception. Ionic conditions play a pivotal role in their structures and functions. To examine the effect of salt ions more systematically, we performed bulk FRET assays for iM molecules with various concentrations of monovalent cations. This study is reminiscent of our previous work, but we focus on a lower pH range and compare this result with the result from hairpin DNA in order to extract features characteristic to iM. In our previous work, we found that Li^+^ destabilizes the iM structure but other monovalent cations stabilize iM either moderately or marginally from smFRET measurements.^17^ Although monovalent cations are known to help stabilize dsDNA, a recent study raised a possibility that they may generally destabilize iM^17,18^. The effect of the type of salt on the stability of iM has not been examined thoroughly or well established yet, so we revisit this problem and aim at gaining a more consistent picture about the effect of salt on iM. As shown in Figure 3a, we found that the melting temperature of iM decreases as the Li^+^ concentration increases, consistent with our previous smFRET work. On the other hand, Na^+^ shows little change in the melting temperature as the concentration increases, and in the case of K^+^, the melting temperature slightly increases as the concentration increases. This result can also be understood in terms of the free energy near the melting temperature (Figure S5). We also performed similar experiments with the same buffer for hairpin DNA as a control sample against iM DNA (Figure 1d, 3e). For all kinds of salts tested, the melting temperature increased by about 20 degrees when the concentration was increased from 50 to 500 mM. The increase in the melting temperature of hairpin DNA is related to the flexibility of ssDNA and enhanced stability of base pairs. ssDNA at high cation concentrations is advantageous for folding into DNA hairpin because the electrostatic repulsion between backbones is reduced by charge screening and ssDNA becomes more flexible. The screening effect would be similar for iM DNA, but the melting temperature of iM did not increase much in the cases of Na^+^ and K^+^, but rather decreased for Li^+^. Therefore, it is reasonable to say that not only Li^+^ but Na^+^ and K^+^ have the considerable destabilizing effect on the iM structure, and this view is consistent with recent studies by other groups.^18^ The mechanism that elucidates the cation-induced instability of the iM structure has not been provided yet. It seems likely that cations in general will make the iM structure less stable by intervening proton-mediated C:C^+^ base pairing which is pivotal for the iM formation. As noted, we showed from smFRET measurements that a high concentration of Li^+^ does not prevent the formation of iM structure, but destabilizes already formed iM structures.^17^ Therefore, the destabilization of iM was related to the size of monovalent cation in solution and augmented in the order of K^+^, Na^+^, and Li^+^, as observed in our experiment. Now we shall interpret the results as follows: Na^+^ and K^+^ have almost no change in melting temperature because the cation (size)-dependent destabilization of iM is largely canceled out or even overturned by the screening-based stabilization of folded nucleic acids. In the case of Li^+^, the size effect is more dominant than the screening effect, so the melting temperature is reduced. We also performed CD measurements, and in contrast to the bulk FRET results, the melting temperature of iM was lowered not only for Li^+^ but for Na^+^ and K^+^ when the salt concentration was increased (Figure S6). It can be speculated that this discrepancy likely originates from the fact that those methods in fact measure different aspects of molecules, FRET measuring the distance between the two positions in the iM molecule and CD measuring the local helicity of the molecule. A recent study indicated that other types of higher-order structure such as C-hairpin, rather than iM, can be formed under high [K^+^] conditions, which may elucidate the current discrepancy because the transition from iM to C-hairpin likely yields the same FRET efficiency maintaining the same inter-dye distance while the helicity specific to iM is disrupted.^19^ Perhaps, this discrepancy may imply that in the presence of Na^+^ and K^+^, some characteristic helical features of iM shall be disrupted by high concentrations of these ions but thanks to significantly reduced charge repulsion, a folded or compact conformation is still maintained, keeping the two (dye-labeled) positions geometrically close. We conclude from both bulk FRET and CD assays that monovalent cations have the destabilizing effect on the iM structure aside from electrostatic stabilization, the degree of which increases in the order of K^+^, Na^+^, and Li^+^.

**Figure 3.**
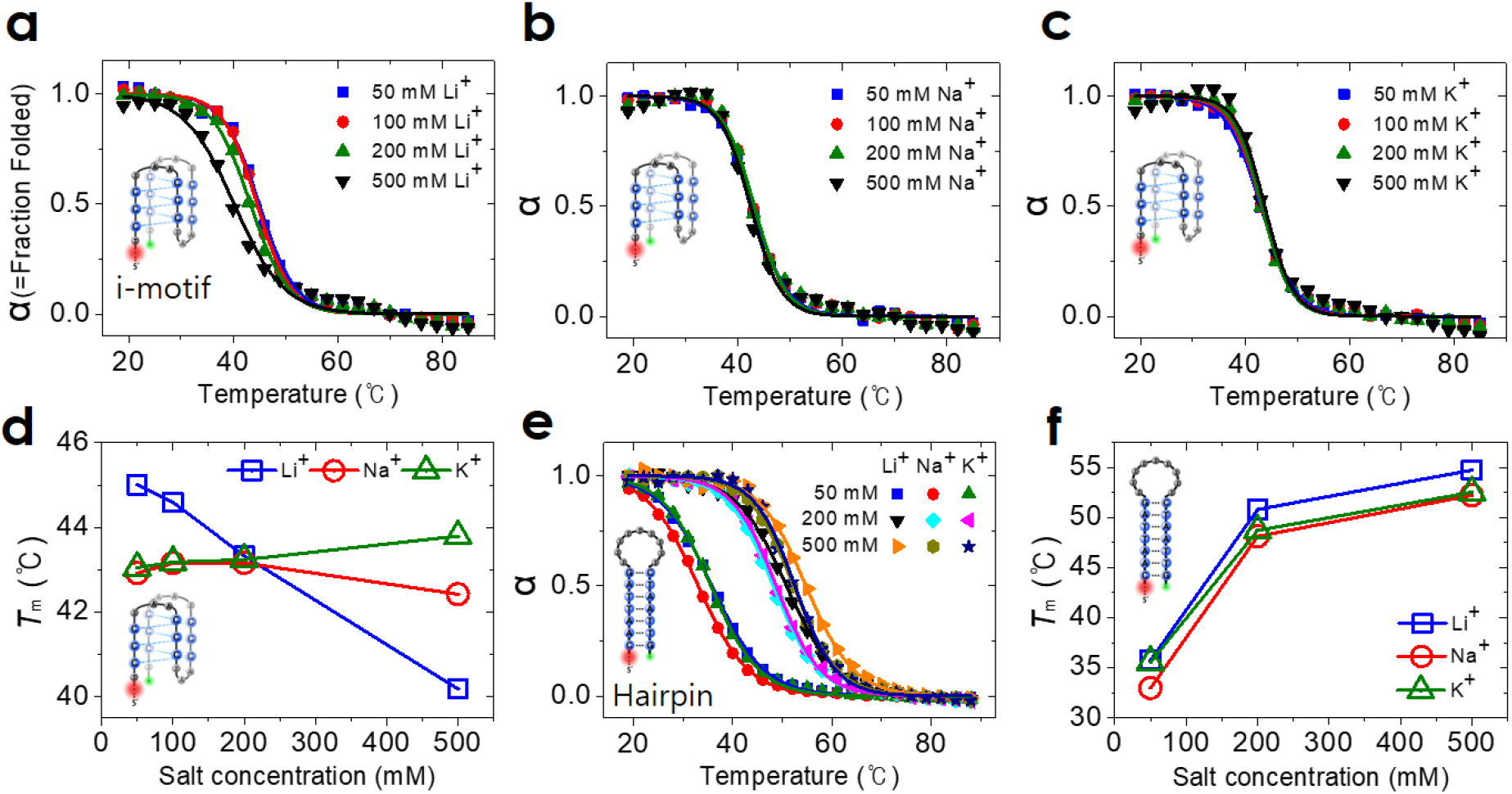
Effect of salt concentration on the melting temperature of (a-d) iM and (e-f) hairpin DNA revealed by bulk FRET assay. (a-c) Displayed are melting curves of iM by C33 at pH = 5.5 in the presence of different cations ((a) Li^+^, (b) Na^+^, (c) K^+^). (d) Melting temperature of iM structures by C32, C33, and C44 at pH 5.5 with various concentrations of different monovalent cations (Li^+^, Na^+^, K^+^). (e) Melting curves of DNA hairpin at pH = 5.5 in the presence of different cations. (f) Melting temperature of DNA hairpin at pH 5.5 with various concentrations of the same monovalent cations.

### Salt-concentration-dependent flexibility of ssDNA

The melting temperature is also related to the flexibility of ssDNA. Flexible ssDNA would be advantageous for iM folding. Cations would make ssDNA as well as dsDNA more flexible by the screening effect. We performed smFRET assays for C33 molecules at pH 6.2 and [Na^+^] = 200 mM. The middle-FRET peak was gradually shifted towards a higher FRET value with increasing concentration of cation, indicating the increased flexibility of ssDNA (Figure 4a). As the concentration of cation increases, the electrostatic repulsion between negatively charged DNA backbones decreases, permitting ssDNA to bend easily and distal ssDNA segments to come closer for compaction. The middle-FRET peak of iM molecules was shifted similarly with increasing concentrations of both Na^+^ and K^+^ (Figure 4a). The middle-FRET peak of iM molecules was shifted more significantly towards high FRET (> 0.5) with increasing concentration of Li^+^, suggesting that the screening effect by Li^+^ is considerably greater than by the other two cations and makes ssDNA more flexible. For instance, the middle-FRET state shifts towards higher FRET value for all three cations tested, but the degree of the shift decreases notably with increasing size of cation (Li^+^<Na^+^<K^+^) at high cation concentration (say, 500 mM) (Figure 4b). We estimated the persistence length of ssDNA under various ionic conditions based on smFRET measurements using the approximate equation 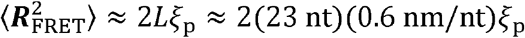 where ***R***_FRET_ is the distance between the donor and acceptor dyes evaluated by FRET, *L* is the contour length of ssDNA between the dyes, and *ξ*_p_ is the persistence length of ssDNA.^30^ From our experimental results and past studies, the persistence length of ssDNA decreases with increasing Na^+^ concentration (Figure 4c).^31^ This is because counterions in buffer would be attracted to the negatively charged DNA backbone, thereby reducing the charge repulsion of the DNA polymer. Studies on the ion atmosphere around DNA have been conducted mainly on duplexes. Small cations can be trapped in the major and minor grooves of DNA and distributed there at high concentrations.^32–34^ Large cations, on the other hand, approach DNA less closely avoiding overlapping with each other, thus reducing occupancy around DNA.^35^ A recent experimental result showed that Li^+^ ions have an advantage over larger cations in occupying the ionic atmosphere around DNA.^36^ Several previous studies confirmed that Li^+^ surrounds double-stranded DNA at high concentrations, but there is no such a study for ssDNA to our knowledge. We propose that smaller cations would also form dense ionic atmosphere around ssDNA. ssDNA is very flexible with the persistence length of ~2 nm, and would be a random, shrunken coil. As in the case of the duplex, smaller cations might be more strongly associated with ssDNA in number and proximity, making ssDNA more flexible. As a result, the smaller the size of cation is in solution, the shorter the persistence length of ssDNA therein would be, which is consistent with our experimental results (Figure 4d). This finding suggests that Li^+^ be advantageous for folding of i-motif compared to other ions for the higher flexibility of the ssDNA chain to be folded (resulting from better screening between contiguous segments of a strand) and the reduced charge repulsion between combined strands (resulting from better screening between distant segments of a strand). In the case of hairpin DNA, which exhibits almost no pH dependence in melting, the melting temperature is about 3 degrees higher with Li^+^ than with Na^+^ (Figure 3f). However, Li^+^ at higher concentrations acts to destabilize i-motif by destabilizing the formation of C:C^+^, and at about 200 mM or higher, the melting temperature of iM is lower with Li^+^ than with other cations (Figure 3d).^17^ In summary, interestingly, prior to iM formation, Li^+^ acts towards folding, but at the moment of folding or after that, it switches to counteract the formation of iM.

**Figure 4.**
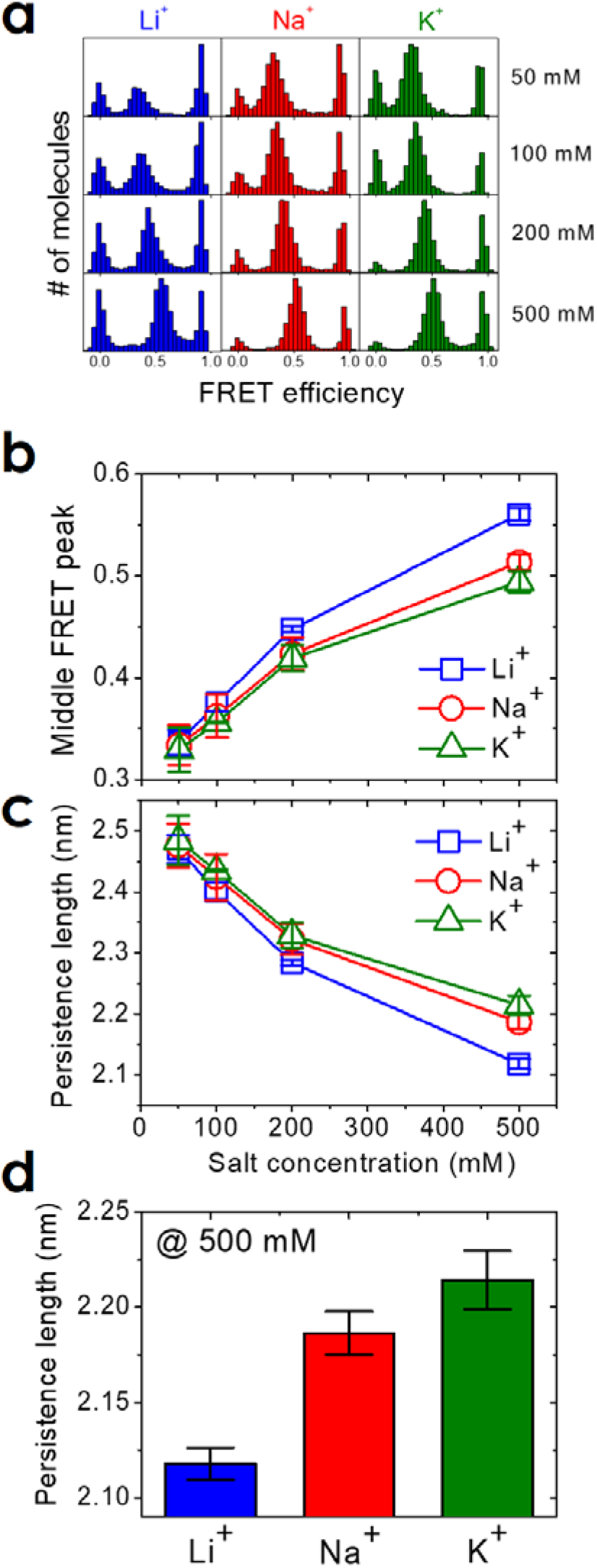
Effect of salt concentration on unfolded single-stranded DNA (C33) at pH 6.2 (a) smFRET histograms of C33 molecules acquired with the indicated concentrations of three different cations. (b) FRET efficiency value of the middle FRET peak increases with salt concentration. (c,d) Screening effect according to the concentration and type of monovalent cation. The persistence length of ssDNA increases in the order of Li^+^, Na^+^, and K^+^. The persistence length of ssDNA at the highest salt concentration used is compared in (d). Error bars represent the sample standard deviation of the corresponding data: Each kind of data is composed of 2 to 4 independent sets of experiments and each set covers hundreds of different DNA molecules.

## IV. CONCLUSIONS

We studied the stability of the i-motif structure formed by three different C-rich sequences (C32, C33, and C44) using FRET methods. As the number of C:C^+^ base pairs increases, the folding free energy decreases linearly to form a more stable iM structure. In addition, we found that the folding free energy per C:C^+^ base pair at room temperature has a lower value for a lower pH, the trend of which is almost linear.

We also investigated the effect of various monovalent cations (Li^+^, Na^+^, K^+^) on the stability of the iM structure. Different from DNA hairpin, model duplex, the iM structure became unstable as the concentration of monovalent cations increased and the destabilization was augmented in the order of K^+^, Na^+^, and Li^+^. In our previous work, Li^+^ was suggested to destabilize the iM structure by competing with H^+^ and therefore weakening the C:C^+^ base pair. On the other hand, at a concentration of cation below 200 mM, the formation of iM was promoted in the order of K^+^, Na^+^, and Li^+^. This is correlated with our observation that the single-stranded chain of an unfolded iM molecule is more flexible and forms a more compact coil in the order of K^+^, Na^+^, and Li^+^. In other words, our finding indicates that smaller cations stick in ssDNA more closely and form a dense cation atmosphere around it, and the screening effect by small cations at high concentration makes ssDNA flexible, which is advantageous for iM formation. The most palpable case here is that of Li^+^: Li^+^ facilitates iM formation by making ssDNA flexible, but makes the iM structure unstable after iM formation.

Our finding in this paper would enhance fundamental understanding about the formation and stability of iM structure and provide insights into the development of iM-based nano-devices.

## AUTHOR INFORMATION

### Notes

The authors declare no competing financial interests.

## ACKNOWLEDGMENT

This work was supported by the Institute for Basic Science (IBS-R023-D1). This work was also partly supported by an NRF grant (2022R1A2B5B01002343) and the Global Research and Development Center Program (2018K1A4A3A01064272) through the NRF of Korea.

## Notes

### Competing Interest Statement

The authors have declared no competing interest.

### Summary of Updates

Corrected due to author order error. (The rest of the text is the same.)

## REFERENCES

(1) Blackburn, E. H. Structure and function of telomeres. Nature 1991, 350, 569–573.

(2) Bryan, T. M.; Cech, T. R. Telomerase and the maintenance of chromosome ends. Curr. Opin. Cell Biol. 1999, 11, 318–324.

(3) Gehring, K.; Leroy, J.; Guéron, M. A tetrameric DNA structure with protonated cytosine. cytosine base pairs. Nature 1993, 363, 561.

(4) Guéron, M.; Leroy, J.-L. The i-motif in nucleic acids. Curr. Opin. Struct. Biol. 2000, 10, 326–331.

(5) Jin, K. S.; Shin, S. R.; Ahn, B.; Rho, Y.; Kim, S. J.; Ree, M. pH-dependent structures of an i-motif DNA in solution. J. Phys. Chem. B 2009, 113, 1852–1856.

(6) Choi, J.; Kim, S.; Tachikawa, T.; Fujitsuka, M.; Majima, T. pH-induced intramolecular folding dynamics of i-motif DNA. J. Am. Chem. Soc. 2011, 133, 16146–16153.

(7) Cui, J.; Waltman, P.; Le, V. H.; Lewis, E. A. The effect of molecular crowding on the stability of human c-MYC promoter sequence I-motif at neutral pH. Molecules 2013, 18, 12751–12767.

(8) Paul, S.; Hossain, S. S.; Samanta, A. Insights into the folding pathway of a c-MYC-Promoter-based i-Motif DNA in crowded environments at the single-molecule level. J. Phys. Chem. B 2020, 124, 763–770.

(9) Megalathan, A.; Wijesinghe, K. M.; Ranson, L.; Dhakal, S. Single-Molecule Analysis of Nanocircle-Embedded I-Motifs under Crowding. J. Phys. Chem. B 2021, 125, 2193–2201.

(10) Zeraati, M.; Langley, D. B.; Schofield, P.; Moye, A. L.; Rouet, R.; Hughes, W. E.; Bryan, T. M.; Dinger, M. E.; Christ, D. I-motif DNA structures are formed in the nuclei of human cells. Nat. Chem. 2018, 10, 631–637.

(11) King, J. J.; Irving, K. L.; Evans, C. W.; Chikhale, R. V.; Becker, R.; Morris, C. J.; Peña Martinez, C. D.; Schofield, P.; Christ, D.; Hurley, L. H. DNA G-quadruplex and i-motif structure formation is interdependent in human cells. J. Am. Chem. Soc. 2020, 142, 20600–20604.

(12) Laisné, A.; Pompon, D.; Leroy, J.-L. [C_7_GC_4_] _4_ association into supra molecular i-motif structures. Nucleic Acids Res. 2010, 38, 3817–3826.

(13) Modi, S.; Krishnan, Y. A method to map spatiotemporal pH changes inside living cells using a pH-triggered DNA nanoswitch. In DNA Nanotechnology, Springer, 2011; pp 61–77.

(14) Wang, Z.-G.; Elbaz, J.; Willner, I. DNA machines: bipedal walker and stepper. Nano Lett. 2011, 11, 304–309.

(15) Nesterova, I. V.; Nesterov, E. E. Rational design of highly responsive pH sensors based on DNA i-motif. J. Am. Chem. Soc. 2014, 136, 8843–8846.

(16) Abou Assi, H.; Garavís, M.; González, C.; Damha, M. J. i-Motif DNA: structural features and significance to cell biology. Nucleic Acids Res. 2018, 46, 8038–8056.

(17) Kim, S. E.; Lee, I.-B.; Hyeon, C.; Hong, S.-C. Destabilization of i-motif by submolar concentrations of a monovalent cation. J. Phys. Chem. B 2014, 118, 4753–4760.

(18) Zhang, F.; Liu, B.; Lopez, A.; Wang, S.; Liu, J. Opposite salt-dependent stability of i-motif and duplex reflected in a single DNA hairpin nanomachine. Nanotechnology 2020, 31, 195503.

(19) Gao, B.; Hou, X.-M. Opposite effects of potassium ions on the thermal stability of i-motif DNA in different buffer systems. ACS omega 2021, 6, 8976–8985.

(20) Goldberg, R. N.; Kishore, N.; Lennen, R. M. Thermodynamic quantities for the ionization reactions of buffers. J. Phys. Chem. Ref. Data 2002, 31, 231–370.

(21) Samuelsen, L.; Holm, R.; Lathuile, A.; Schönbeck, C. Buffer solutions in drug formulation and processing: How pKa values depend on temperature, pressure and ionic strength. Int. J. Pharm. 2019, 560, 357–364.

(22) Kim, S.-E.; Lee, I.-B.; Hong, S.-C. The effect of the oxygen scavenging system on the pH of buffered sample solutions: in the context of single-molecule fluorescence measurements. Bull. Korean Chem. Soc. 2012, 33, 958–962.

(23) Kang, C.; Berger, I.; Lockshin, C.; Ratliff, R.; Moyzis, R.; Rich, A. Stable loop in the crystal structure of the intercalated four-stranded cytosine-rich metazoan telomere. Proceedings of the National Academy of Sciences 1995, 92, 3874–3878.

(24) Catasti, P.; Chen, X.; Deaven, L. L.; Moyzis, R. K.; Bradbury, E. M.; Gupta, G. Cystosine-rich strands of the insulin minisatellite adopt hairpins with intercalated Cytosine+· Cytosine pairs. J. Mol. Biol. 1997, 272, 369–382.

(25) Phan, A. T.; Leroy, J.-L. Intramolecular i-motif structures of telomeric DNA. J. Biomol. Struct. Dyn. 2000, 17, 245–251.

(26) Mathur, V.; Verma, A.; Maiti, S.; Chowdhury, S. Thermodynamics of i-tetraplex formation in the nuclease hypersensitive element of human c-myc promoter. Biochem. Biophys. Res. Commun. 2004, 320, 1220–1227.

(27) Xu, Y.; Sugiyama, H. Formation of the G-quadruplex and i-motif structures in retinoblastoma susceptibility genes (Rb). Nucleic Acids Res. 2006, 34, 949–954.

(28) Khan, N.; Aviñó, A.; Tauler, R.; González, C.; Eritja, R.; Gargallo, R. Solution equilibria of the i-motif-forming region upstream of the B-cell lymphoma-2 P1 promoter. Biochimie 2007, 89, 1562–1572.

(29) Školáková, P.; Renčiuk, D.; Palacký, J.; Krafčík, D.; Dvořáková, Z.; Kejnovská, I.; Bednářová, K.; Vorlíčková, M. Systematic investigation of sequence requirements for DNA i-motif formation. Nucleic Acids Res. 2019, 47, 2177–2189.

(30) Phillips, R.; Kondev, J.; Theriot, J.; Garcia, H. Physical biology of the cell; Garland Science, 2012.

(31) Murphy, M.; Rasnik, I.; Cheng, W.; Lohman, T. M.; Ha, T. Probing single-stranded DNA conformational flexibility using fluorescence spectroscopy. Biophys. J. 2004, 86, 2530–2537.

(32) Chu, V. B.; Bai, Y.; Lipfert, J.; Herschlag, D.; Doniach, S. Evaluation of ion binding to DNA duplexes using a size-modified Poisson-Boltzmann theory. Biophys. J. 2007, 93, 3202–3209.

(33) Yoo, J.; Aksimentiev, A. Competitive binding of cations to duplex DNA revealed through molecular dynamics simulations. J. Phys. Chem. B 2012, 116, 12946–12954.

(34) Giambaşu, G. M.; Gebala, M. K.; Panteva, M. T.; Luchko, T.; Case, D. A.; York, D. M. Competitive interaction of monovalent cations with DNA from 3D-RISM. Nucleic Acids Res. 2015, 43, 8405–8415.

(35) Bai, Y.; Greenfeld, M.; Travers, K. J.; Chu, V. B.; Lipfert, J.; Doniach, S.; Herschlag, D. Quantitative and comprehensive decomposition of the ion atmosphere around nucleic acids. J. Am. Chem. Soc. 2007, 129, 14981–14988.

(36) Gebala, M.; Bonilla, S.; Bisaria, N.; Herschlag, D. Does cation size affect occupancy and electrostatic screening of the nucleic acid ion atmosphere? J. Am. Chem. Soc. 2016, 138, 10925–10934.

