## supplementary material for "Two opposing effects of monovalent cations on the stability of i-motif structure"

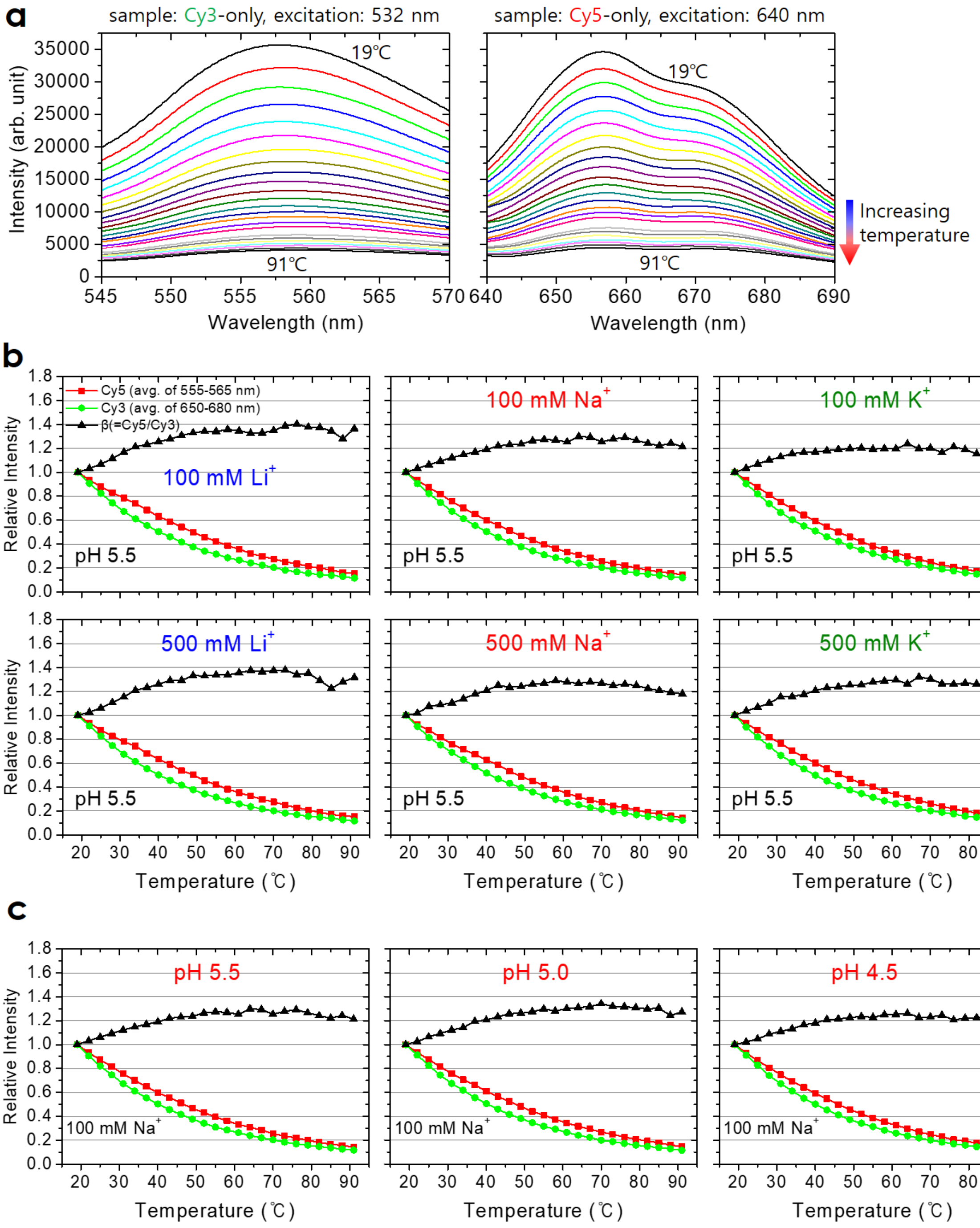


**Figure S1. Effect of temperature on the fluorescence intensity of Cy3 and Cy5 dyes. This result is used to adjust the bulk FRET data (the detailed procedure of calibration is shown in Figure S3). We used a sequence for each dye: 5'-(Cy3)-TTTTT for Cy3 and 5'-(Cy5)-TTTTT for Cy5. (a) Fluorescence intensities of Cy3 and Cy5 dyes with increasing temperature in 100 mM $Na^+$ at pH 5.5. As the temperature increases, the fluorescence intensity decreases. (b) The fluorescence intensities of Cy3 and Cy5 decrease with the temperature under various salt conditions. These fluorescent dyes exhibit almost the same temperature dependence in the two different concentrations (100 mM and 500 mM) of a given salt. Therefore, we adjusted raw experimental data acquired with different concentrations (50, 100, 200, and 500 mM) of a salt with the dye-**

**calibration fluorescence data (dye-only data) acquired with 100 mM solution of the same type of salt. (c) The fluorescence intensities of Cy3 and Cy5 in 100 mM $Na^+$ at various pH values decrease with the temperature. There is no obvious pH dependence of fluorescence intensities. Panel (b) and (c) show three relative intensities of dye fluorescence. The intensity of Cy5 fluorescence normalized by the intensity at 19$^o$C, the intensity of Cy3 fluorescence normalized by the intensity at 19$^o$C, and the ratio (β) of normalized Cy5 fluorescence to normalized Cy3 fluorescence are shown in red, green, and black, respectively.**

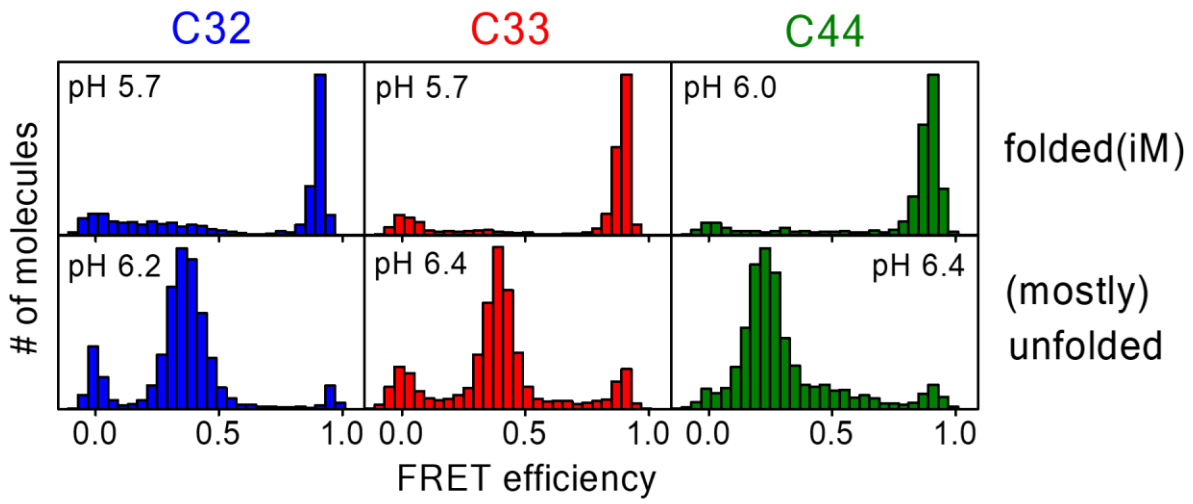


**Figure S2. FRET efficiency histograms of C32, C33, and C44 molecules in 200 mM $Na^+$ at the minimum and maximum pH values tested. For each molecule, the molecule appears mainly folded at the minimum pH values (top row) and mainly unfolded at the maximum pH values (bottom row).**

**How to obtain the fraction folded (α) in the bulk-FRET-based melting assay.**

Since the fluorescence intensities of Cy3 and Cy5 dyes measured from Cy3-only and Cy5-only samples, respectively, decrease with the temperature (Figure S1), this temperature dependence needs to be compensated for to identify the intensity change due to FRET only. A series of normalization, correction, and rescaling steps are explained and illustrated in Figure S3 and its legend. In (c), the ratio of fluorescence intensities appears to reach a maximum simply because the temperature dependent decrease of the fluorescence intensity of Cy3 is more rapid than that of Cy5, so this part becomes flat after correcting the effect. The melting temperature ($T_m$) is defined as the midpoint of the transition.

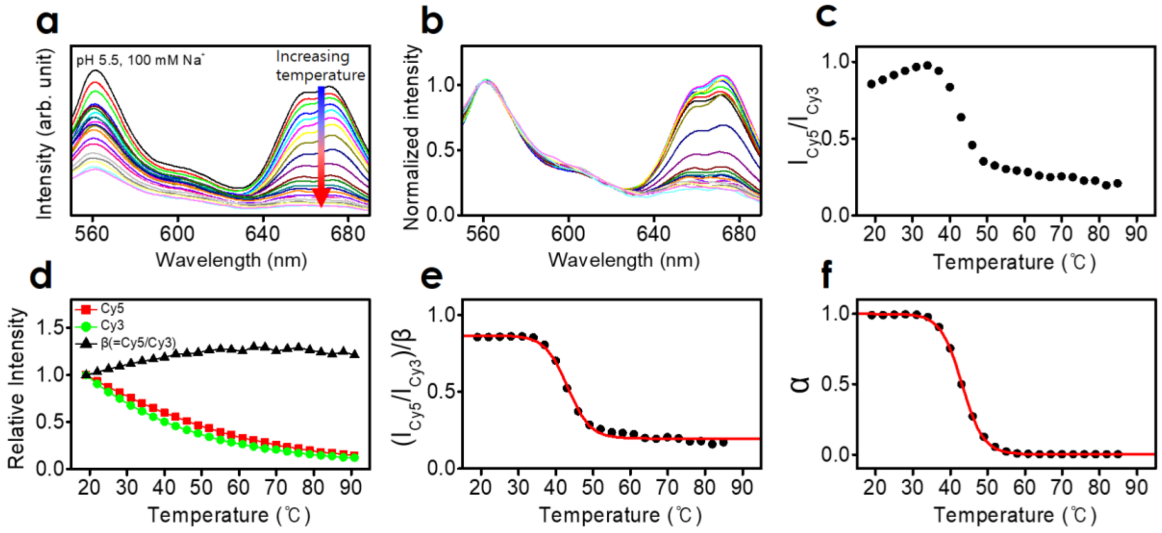


**Figure S3. Procedure to obtain the fraction folded(=α). (a) Raw emission spectra of the C33 sample excited at λ = 532 nm. The curves in different colors show the spectra taken at different temperatures as indicated by a red arrow. (b) The fluorescence intensity of each curve is normalized by the fluorescence intensity of the same curve averaged over the range of 555 - 565 nm. After normalization, the fluorescence intensity at 560 nm becomes 1. (c) Temperature-dependent ratio of fluorescence intensity of Cy5 to that of Cy3, which is given by the fluorescence intensity averaged over the range of 650-680 nm from Panel (b) acquired at various temperatures. In Panel (b), the fluorescence intensity of Cy3 is already normalized to 1. (d) The fluorescence intensities of dye-only samples and their ratio β (see Figure S1) with respect to the sample temperature. Cy3-only and Cy5-only samples are excited at λ = 532 nm and 640 nm, respectively. The intensity values are normalized by the maximum value acquired, the intensity at 19°C. The ratio is used as a correction factor for the relative intensity (see Figure S1). (e) The ratio shown in (c) is adjusted by β and fit with a sigmoidal function. (f) The ratio curve in (e) is rescaled to set the values at 5 °C and 95 °C to α = 1 and 0, respectively.**

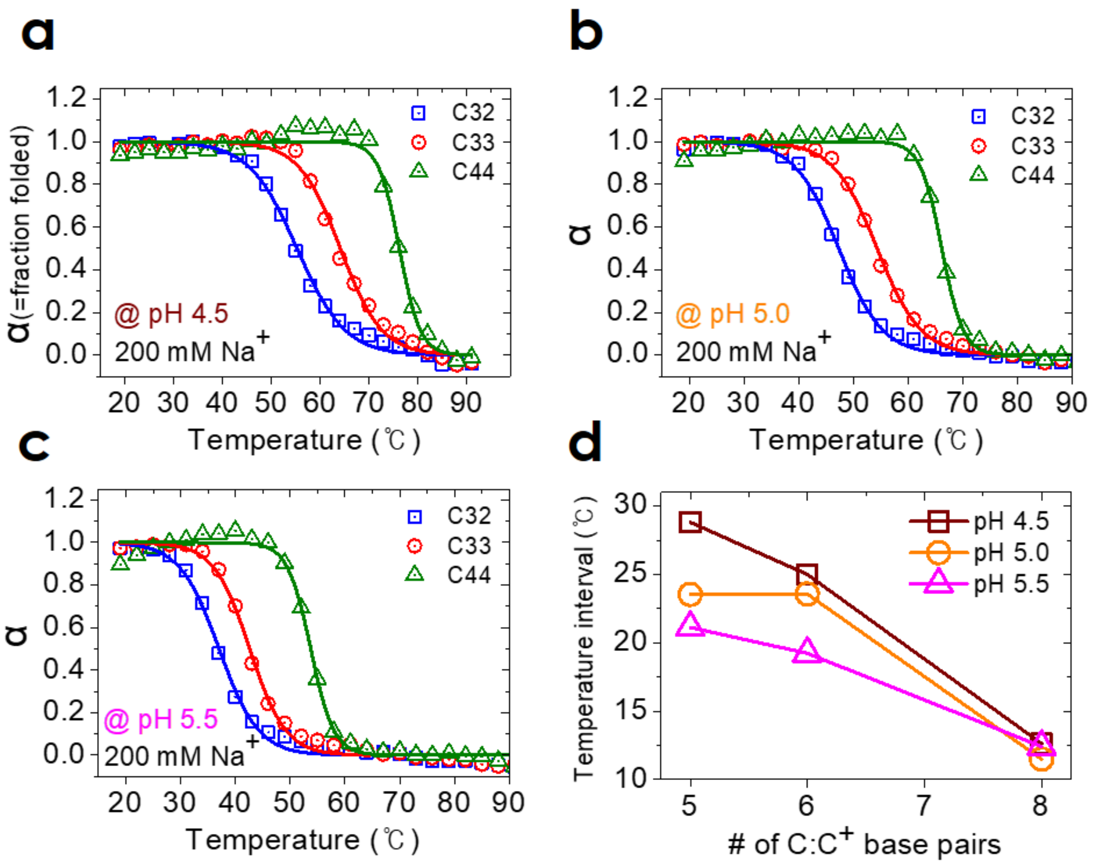


**Figure 4. (a-c) Melting curves of iM samples at various pH values. The melting curves at pH 5.5 are shown in the main text and also shown here for easy comparison. (d) Temperature interval of the melting transition with respect to the number of C:C+ pairs. The temperature interval is defined by the difference of the temperatures for α = 5% and 95%.**

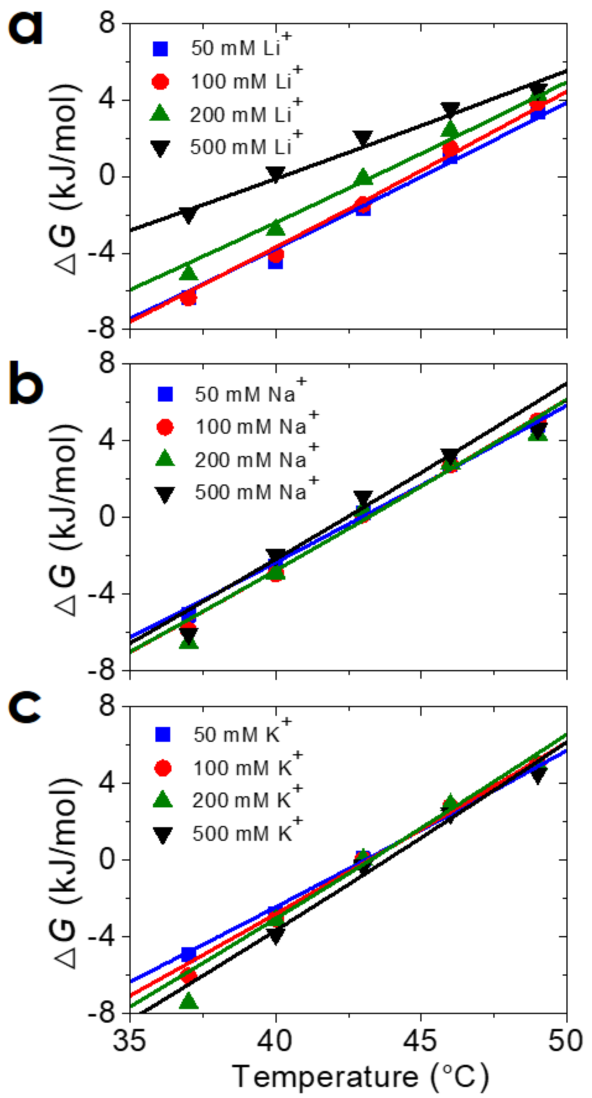


**Figure S5. Folding free energy ($\Delta G$ = $RT$ ln$K_{eq}$ where $K_{eq}$ = [folded]/[unfolded] = α/(1-α)) of C33 DNA at various salt concentrations.**

**Circular Dichroism (CD) assay**

Circular dichroism spectra were acquired using a Jasco-815 spectropolarimeter (Jasco, Japan) equipped with a Peltier temperature controller. We acquired CD spectra as follows: iM molecules were loaded in a standard quartz cuvette and their spectra were acquired in the range of 240 - 320 nm with the scanning interval and speed of 0.5 nm and 100 nm/min, respectively. Each spectrum was obtained by subtracting a buffer-only spectrum from a raw sample spectrum. A final CD spectrum was acquired by averaging three runs of measurement (Figure S6a).

The DNA oligonucleotide used in CD measurements was 5'-TCC CTA ACC CTA ACC CTA ACC CT-3', the same C-rich human telomere sequence as those used in FRET assays, and purchased from Bioneer (Daejeon, Korea). The oligo sample was dissolved to a final concentration of 4.4 μM (equiv. OD = 0.9) in 20 mM sodium acetate buffer (pH 5.5). The melting assay via CD measurement was carried out by increasing the temperature gradually from 25°C (in 5°C increments) and measuring spectra at each target temperature after reaching it and staying there stably for 4 minutes. The peak wavelength $\lambda_{\mathrm{peak}}^{\mathrm{CD}}$ was found by a parabolic fit to the peak region of the CD curve (within ±10 nm from an estimated peak position) (Figure S6b). The melting temperature $T_{\mathrm{m}}$ was obtained by fitting a sigmoidal function to the $\lambda_{\mathrm{peak}}^{\mathrm{CD}}$-vs.-temperature graph (Figure S6 c,d).

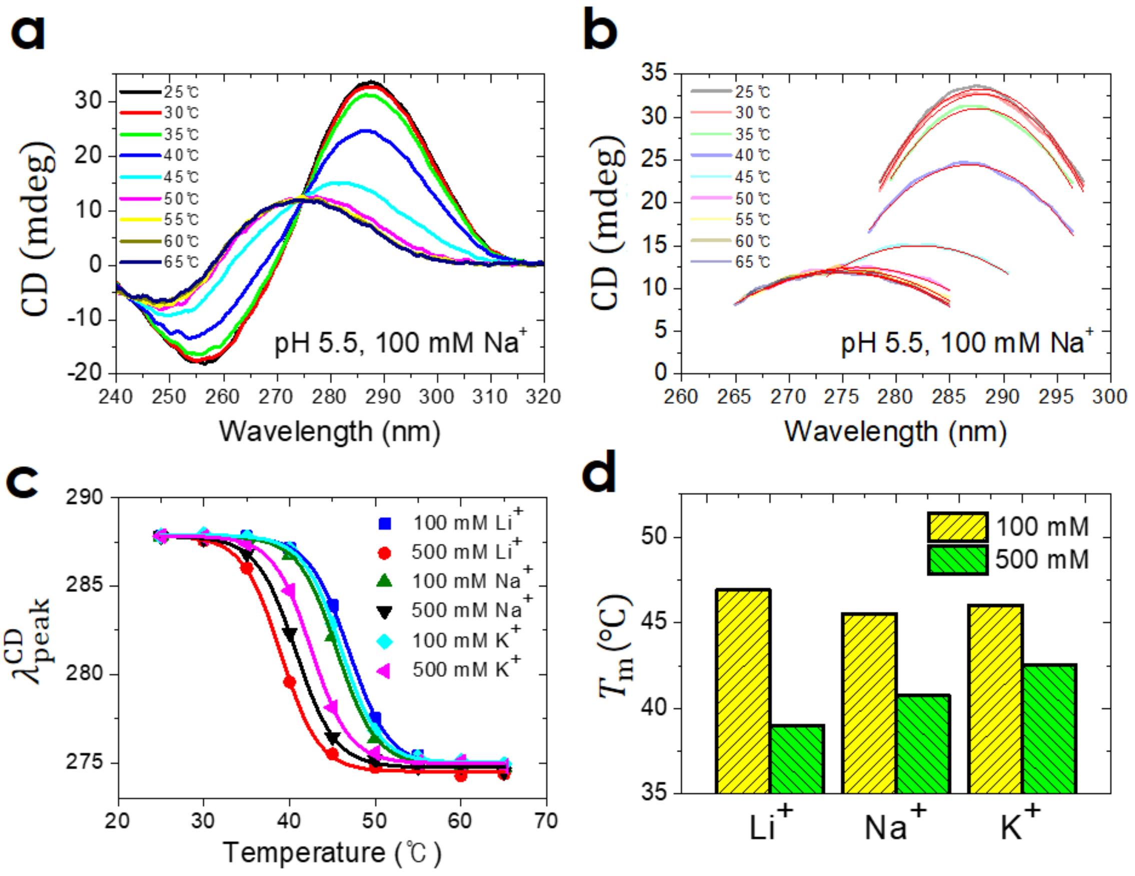


**Figure S6. (a) Representative data of CD spectra (100 mM $Na^{+}$) (b) Parabolic fit to the peak area (±10 nm with respect to the peak position) of the CD curve (c) $\lambda_{peak}^{CD}$-vs.-temperature graph for various salt conditions (d) Melting temperature of iM from CD spectra according to the type and concentration of salt (obtained from Figure S6c).**
